# Scallop enables accurate assembly of transcripts through phasing-preserving graph decomposition

**DOI:** 10.1101/123612

**Authors:** Mingfu Shao, Carl Kingsford

## Abstract

We introduce Scallop, an accurate, reference-based transcript assembler for RNA-seq data. Scallop significantly improves reconstruction of multi-exon and lowly expressed transcripts. On 10 human samples aligned with STAR, Scallop produces (on average) 35.7% and 37.5% more correct multi-exon transcripts than two leading transcript assemblers, StringTie [1] and TransComb [2], respectively. For transcripts expressed at low levels in the same samples, Scallop assembles 65.2% and 50.2% more correct multi-exon transcripts than StringTie and TransComb, respectively. Scallop obtains this improvement through a novel algorithm that we prove preserves all phasing paths from reads (including paired-end reads), while also producing a parsimonious set of transcripts and minimizing coverage deviation.

---

RNA sequencing (RNA-seq) is an established technology that enables identification of novel genes and splice variants as well as accurate measurement of expression abundances [3]. The RNA-seq protocol produces short sequencing reads sampled from the expressed transcripts, and transcript assembly is the fundamental computational problem to reconstruct the full-length expressed transcripts from the reads. This step is crucial for transcript quantification and differential expression analysis, and also plays a central role in revealing tissue-specific splicing patterns [4] and understanding the regulation of gene expressions [5].

Transcript assembly methods can be divided into reference-based (or genome-guided) methods and *de novo* methods, depending on whether a reference genome is assumed to be available. Reference-based methods (e.g., Cufflinks [6], Scripture [7], IsoLasso [8], SLIDE [9], CLIIQ [10], CEM [11], MITIE [12], iReckon [13], Traph [14], Bayesembler [15], StringTie [1], CIDANE [16] and TransComb [2]) usually first use the read alignments produced by an RNA-seq aligner (e.g., TopHat2 [17], SpliceMap [18], STAR [19], and HISAT [20]) to build a so-called splice graph for each gene loci. In the splice graph, vertices correspond to exons (or partial exons), edges correspond to splice junctions, and coverage of exons and abundance of splice junctions are encoded as weights of vertices or edges. The expressed transcripts, represented as a set of paths of the splice graph, are inferred so as to mostly fit the topology and the weights of the splice graph. For instance, StringTie [1] iteratively computes the heaviest path in the splice graph, collect that path, and updates the weights of the remaining splice graph via a max-flow formulation. TransComb [2] employs a bin-packing strategy to gradually reconstruct paths guided by the weighted junction graph [21]. *De novo* assembly methods (e.g., TransABySS [22], Rnnotator [23], Trinity [24], SOAPdenovo-Trans [25], Velvet [26], Oases [27], IDBA-Tran [28], and BinPacker [21]), are mainly used for non-model species and cancer samples, for which a reference genome is unavailable or significantly diverged. When a high-quality reference genome is available, reference-based methods usually obtain better accuracy, since they can tolerate much lower read coverage inside exons.

Transcript assembly remains an open and challenging problem, due to the ubiquity of paralogs, unevenness of read coverage, and diversity of splice variants. According to the benchmarking studies [29, 30], the accuracies of existing transcript assembly methods are still very low, especially for lowly expressed transcripts and those genes with multiple spliced isoforms. Hence, new algorithmic ideas are needed to produce more accurate transcript assemblies.

Scallop is a reference-based transcript assembler that enables accurate identification of multi-exon transcripts and lowly expressed transcripts. Scallop obtains higher accuracy through a novel algorithm to decompose splice graph into transcripts. Our algorithm fully takes advantage of the phasing information derived from the reads (including paired-end reads) spanning more than two exons. Such phasing information is organized as a set of phasing paths, and our algorithm can be proved to preserve all phasing paths (except for those with false negative edges). This theoretical guarantee is achieved through subroutines to decompose the splice graph so as to not to break any phasing path. In addition, our algorithm simultaneously optimizes two other objectives of minimizing the read coverage deviation and minimizing the number of expressed transcripts. To minimize the deviation from the observed read coverage, we formulate and solve a linear programming problem as a subroutine. We also identify when these linear programming instances have multiple optimal solutions and use the abundance of the phasing paths to adjust them via another linear programming problem. To minimize the number of predicted transcripts following the parsimony principle, we devise an efficient subroutine to reduce an upper bound on the required paths. This subroutine can also naturally identify false positive edges in the splice graph. All three objectives are unified into a single iterative optimization framework, and act together to cause Scallop to possess both high sensitivity (through fully using the phasing information and minimizing the coverage deviation) and high precision (through minimizing the number of reconstructed transcripts and removing false positive edges).

We compare Scallop with two recent and popular reference-based transcript assemblers, StringTie and TransComb. We first evaluate them using 10 human RNA-seq samples, all of which use strand-specific and paired-end protocols (Supplementary Table 2). Among them, ST1, ST2, and ST3 were evaluated in the StringTie paper, TC1 and TC2 were used in the TransComb paper, while the other five (SC1, SC2, …, SC5) were chosen by us from ENCODE project (https://genome.ucsc.edu/ENCODE, 2003–2012). We have tuned the parameters of Scallop on the first 5 samples (i.e., ST1, ST2, ST3, TC1 and TC2, hereinafter we call them training samples) and after that we froze Scallop and tested it on other 5 samples (testing samples).

For each of the 10 samples, we experiment with three RNA-seq aligners, TopHat2, STAR, and HISAT2, to map the sequenced reads to the reference genome. Taking the reads alignment as (the only) input, each method predicts a set of expressed transcripts. (The current version of TransComb does not support HISAT2 alignments, so we only compare StringTie and Scallop when using HISAT2 alignments.) Since we do not have a ground truth set of expressed transcripts, we evaluate the predicted transcripts by comparing them with the entire annotation database of known human transcripts, as is commonly done [15, 1, 16, 2]. Usually for a given sample only a small subset of the transcripts in the database are expressed, and it is also likely that some predicted transcripts are novel and thus are not in the current database. A multi-exon transcript is defined as correct if its exon chain can be exactly matched to a known (multi-exon) transcript, while a single-exon transcript is correct if it overlaps at least 80% with a known single-exon transcript. We use the gffcompare program to determine whether a predicted transcript is correct. Sensitivity is then taken to be the ratio between the number of correct transcripts and the total number of known transcripts, and precision is the ratio between the number of correct transcripts and the total number of predicted transcripts.

All three methods support specifying a parameter, minimum coverage threshold, to control the minimum expression abundance of the predicted transcripts. Since highly expressed transcripts are easier to assemble, this parameter essentially balances the sensitivity and precision of the predicted transcripts. In our experiments, to evaluate the capability of these methods in balancing sensitivity and precision, we run these methods on 10 different thresholds across a reasonable range: {0, 1, 2.5, 5, 7.5, 10, 25, 50, 75, 100}.

The accuracy of these three assemblers using different alignment programs and different minimum coverage thresholds is shown in Figure 1 (for the 5 testing samples) and Supplementary Figure 1 (for the 5 training samples). The trade-off between sensitivity and precision of multi-exon transcripts as the minimum coverage threshold is varied is shown in Figure 1A and Supplementary Figure 1A. The curves for Scallop are highest for all 10 samples and for all three aligners, indicating that no matter the desired sensitivity-precision trade-off, Scallop outperforms the other two assemblers in reconstructing multi-exon transcripts. Accuracy is further summarized as the area under the precision-sensitivity curve (AUC), shown in Figure 1B and Supplementary Figure 1B. With TopHat2 alignments, the average AUC score of Scallop over the 5 testing samples is 22.0% and 21.8% higher than that of StringTie and TransComb, respectively. With STAR alignments, the improvement is 34.1% over StringTie and 106% over TransComb. With HISAT2 alignments, Scallop’s AUC is 23.5% higher than that of StringTie.

**Figure 1:**
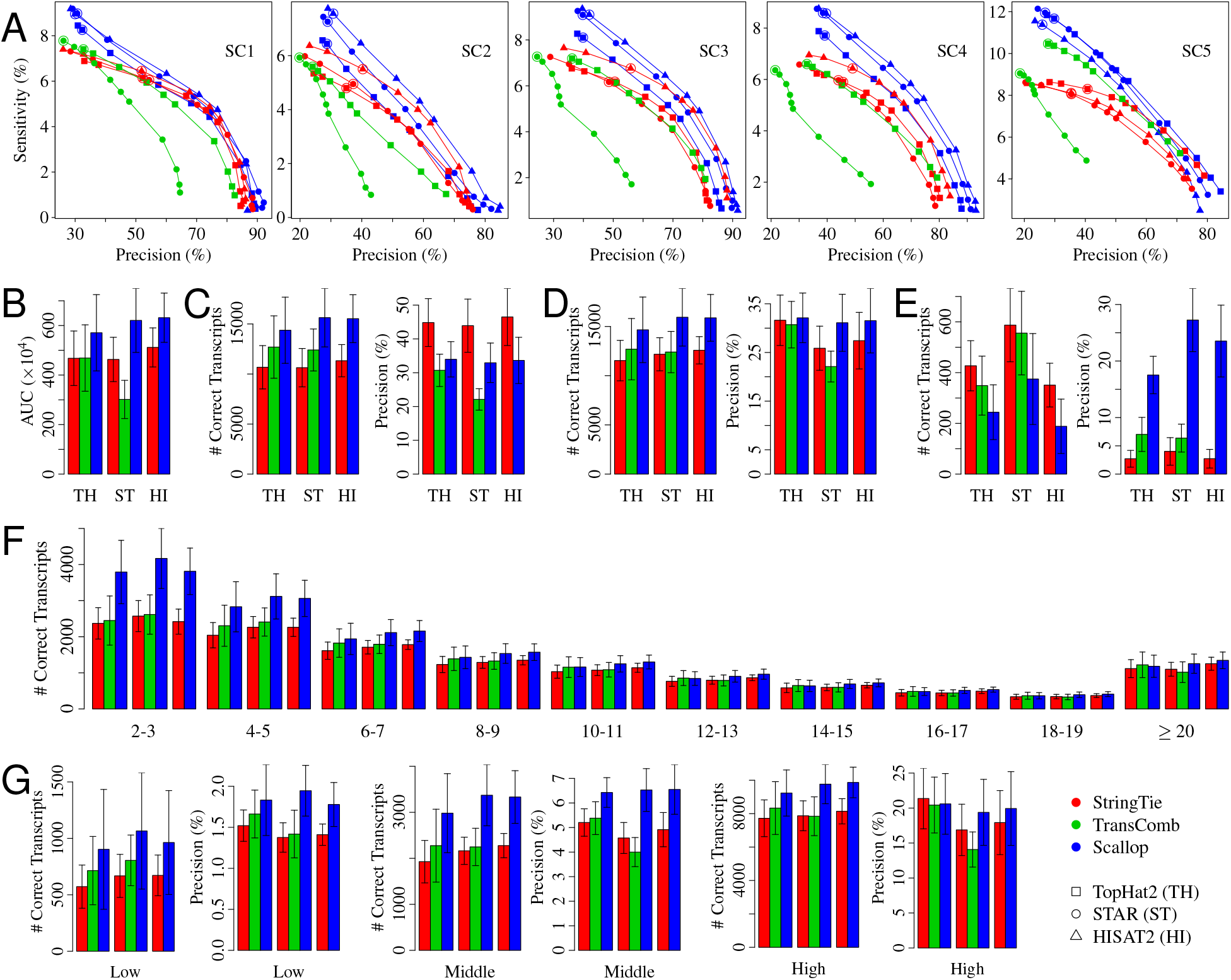
Comparison of the three methods (StringTie, TransComb, and Scallop) over the 5 testing samples. (A) The precision-sensitivity curves for multi-exon transcripts. Each curve connects 10 points, correspond-ing to the 10 different minimum coverage thresholds {0, 1, 2.5, 5, 7.5, 10, 25, 50, 75, 100}; the default value of this parameter is circled. (B) The average AUC (area under the precision-sensitivity curve) over the 5 samples. The three groups of bars correspond to TopHat2, STAR, and HISAT2 alignments, respectively (the same for other panels). The error bars show the standard deviation over the 5 samples (the same for other panels). (C) The average sensitivity and precision of multi-exon transcripts for methods running with default parameters. (D) The average sensitivity and precision of multi-exon transcripts for methods running with minimum coverage set to 0. (E) The average sensitivity and precision of single-exon transcripts for methods running with default parameters. (F) The average number of correct transcripts with different number of exons for methods running with minimum coverage set to 0. (G) The average sensitivity and precision of multi-exon transcripts with each subset of transcripts (corresponding to low, middle, and high expression level) as ground truth for methods running with minimum coverage set to 0.

The default values of the minimum coverage threshold used by StringTie, TransComb and Scallop are 2.5, 0, and 1.0 respectively (circled in Figure 1A and Supplementary Figure 1A). At default parameters, Scallop is significantly more sensitive than StringTie and TransComb at detecting multi-exon transcripts (Figure 1C and Supplementary Figure 1C): averaged over the 5 testing samples, Scallop produces between 34.7%–47.0% more correct multi-exon transcripts than StringTie, and between 13.2%–26.0% more than TransComb, depending on the aligners used. StringTie’s higher precision at default parameters can be explained by its higher default minimum coverage threshold. When evaluated at equivalent sensitivity (Figure 1A and Supplementary Figure 1A), Scallop obtains higher precision than both StringTie and TransComb. In particular, Scallop consistently outperforms StringTie and TransComb in terms of both sensitivity and precision when the minimum coverage threshold is set to 0 (i.e., sensitivity is maximized) for all three methods (Figure 1D and Supplementary Figure 1D). On average over the 5 testing samples, Scallop predicts 26.2%– 30.9% and 15.5%–28.5% more correct multi-exon transcripts than StringTie and TransComb, depending on the aligners used. (A comparison of the sets of correct multi-exon transcripts predicted by three methods is given in Supplementary Figure 2.) Further, at minimum coverage equal to 0, the three methods obtain similar multi-exon precision when using TopHat2 alignments, while for STAR and HISAT2 alignments Scallop obtains a significantly higher precision than both StringTie and TransComb.

StringTie and TransComb obtain higher sensitivity but lower precision than Scallop on single-exon transcripts with default parameters (Figure 1E and Supplementary Figure 1E). However, the overall number of correct single-exon transcripts obtained by these methods is relatively small compared with that of multi-exon transcripts (compare scale of Figure 1E with Figure 1C). Scallop aggressively filters short and lowly expressed single-exon transcripts to make the precision of single-exon transcripts comparable to that of multi-exon transcripts.

We further compare the number of correct transcripts with different numbers of exons when their sensitivity is maximized (i.e., the minimum coverage threshold is 0). The results are shown in Figure 1F and Supplementary Figure 1F. While Scallop is able to identify more correct transcripts for genes with at least up to 17 exons, its advantage is most pronounced on transcripts with 2–7 exons. For example, with TopHat2 alignments, on average over the testing samples, Scallop obtains 60.0% and 54.9% more correct transcripts with 2 or 3 exons than StringTie and TransComb, respectively.

Scallop also significantly improves identification of lowly expressed transcripts. To perform a quantitative measurement, we use Salmon [31] to quantify the 10 RNA-seq samples using human annotation database (ENSEMBL release 87) as reference. For each sample, we collect all multi-exon transcripts with expression abundance larger than a threshold (TPM ≥ 0:1), sort them according to their expression abundances, and divide them into three equal subsets corresponding to low, middle and high expression levels. We then compute the accuracy of the three methods on multi-exon transcripts with each subset as the ground truth (Figure 1G and Supplementary Figure 1G). Scallop achieves higher accuracy on all three expression levels, but the advantage is much more significant for low and middle levels. For example, with STAR alignment, on average over the 5 testing samples, Scallop obtains 59.6%, 55.8%, and 24.1% more correct multi-exon transcripts than StringTie for low, middle, and high expression levels, respectively.

To show the generality of the above results, we collected all human RNA-seq paired-end samples from the ENCODE project (https://www.encodeproject.org, 2013–present) that provide pre-computed read alignments (for experiments with more than one such sample, we arbitrarily select one). This yielded 50 strand-specific and 15 non-strand-specific samples. Since the three methods use different parameters to balance precision and sensitivity, to compare them on equal footing, we compute an adjusted sensitivity and adjusted precision. Specifically, for each sample, we fix the precision φ of the method with highest precision, and for each of the other two methods with smaller precision, we discard its predicted transcripts from the lowest coverage until the precision equals ϕ; the adjusted sensitivity is the sensitivity at this precision ϕ. We compute the adjusted precision analogously, filtering low-coverage transcripts of the two methods with higher sensitivity until all methods have the same sensitivity.

Scallop shows higher adjusted sensitivity and adjusted precision than both StringTie and TransComb on nearly all samples when using default parameters (Supplementary Figure 3 and Supplementary Figure 4). On average over these 50 strand-specific samples, Scallop obtains 17.4% and 19.3% more correct multiexon transcripts (after adjustment to identical precision) than StringTie and TransComb, respectively. The average adjusted precision are 39.1%, 38.6%, and 47.3% for StringTie, TransComb, and Scallop, respectively. On average over the 15 non-strand-specific samples, Scallop obtains 14.6% more correct multi-exon transcripts (after adjustment) than StringTie and obtains an average adjusted precision of 48.0%, while that of StringTie is 42.0%. (TransComb fails on all 15 non-strand-specific samples.) Scallop’s advantage is even more pronounced when lowly expressed transcripts are included by setting the minimum coverage to 0 for all three methods (Supplementary Figure 5 and Supplementary Figure 6). This provides additional evidence that Scallop is particularly more sensitive to lowly expressed transcripts.

Scallop has a comparable but slightly longer running time than StringTie, while TransComb takes significantly longer (Supplementary Figure 7). On average over the 10 samples and the 10 minimum coverage thresholds, with TopHat2 alignments, the running times of Scallop and TransComb are 1.08 × and 3.78 × that of StringTie. With STAR alignments, Scallop and TransComb take 1.34 × and 5.42 × longer than StringTie, respectively. With HISAT2 alignments, Scallop takes 1.25 × longer than StringTie.

Scallop presents a new technique for estimating transcriptome assembly from RNA-seq. While building upon the standard paradigm of the splice graph, it uses a novel algorithm to decompose the graph through optimizing several competing objectives. This leads it to achieve both higher sensitivity and higher precision over a wide range of minimum coverage thresholds. In particular, Scallop can theoretically guarantee to fully use the phasing information to resolve complicated alternative splicing variants, causing significant improvement on assembling multi-exon transcripts and lowly expressed transcripts. Scallop is freely available as open source at http://www.github.com/Kingsford-Group/scallop.

## Acknowledgements

This research is funded in part by the Gordon and Betty Moore Foundations Data-Driven Discovery Initiative through Grant GBMF4554 to C.K., by the US National Science Foundation (CCF-1256087, CCF-1319998) and by the US National Institutes of Health (R01HG007104).

## Online Methods

### Problem Statement

Based on a given alignment of sequencing reads to a reference genome, we build the splice graph, denoted as *G* = (*V, E*), as follows. We first extract splice positions from the alignments. These splice positions imply the boundaries of exons (or partial exons) and introns of the reference genome. For each inferred exon, we add a vertex *v* to *V*. If there exist reads spanning two exons *u* and *v* (where *u* occurs before *v* in the genome), we add a directed edge *e* = (*u, v*) to *E*, and set the weight of *e*, denoted as *w*(*e*), to the number of such reads that span *u* and *v*. We also add a source vertex *s*, and for each vertex *u* ∈ *V* \ {*s*} with in-degree of 0, we add a directed edge (*s, u*) with weight 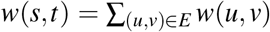. Similarly, we add a sink vertex *t*, and for each vertex *v* ∈ *V* \ {*s*,*t*} with out-degree of 0, we add a directed edge (*v*,*t*) to *E* with weight 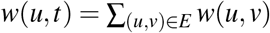. See Figure 2 for an example.

**Figure 2:**
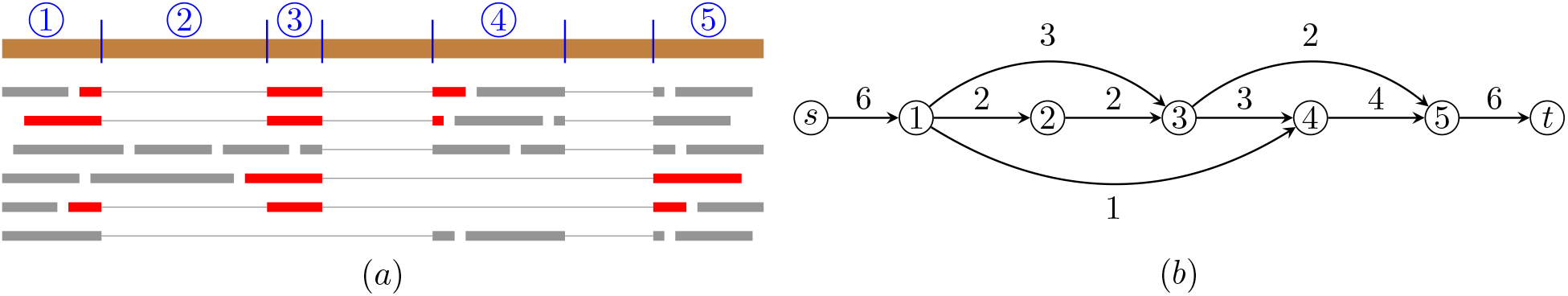
Example of building splice graph and phasing paths. (**a**) Alignment of reads to the reference genome. Inferred (partial) exons are marked with blue numbers. Reads that span more than two exons are marked red, from which we can get the set of phasing paths as {(1, 3, 4), (2, 3, 5), (1, 3, 5)}. The abundance of these phasing paths are *g*(1, 3, 4) = 2, *g*(2, 3, 5) = 1, and *g*(1, 3, 5) = 1. (**b**) The corresponding splice graph and weights for all edges.

Many reads (including paired mates) can span more than two exons, providing phasing information to reconstruct the expressed transcripts. We collect such phasing information as a set of phasing paths of *G*, denoted as *H*, as follows. If a read spans vertices 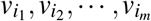 of *G* (i.e., this read sequentially aligns to these corresponding exons), *m* ≥ 3, we then add a phasing path 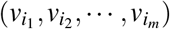 to *H*. For the case of paired-end reads, if we have that one read span vertices 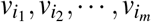, and its mate read spans vertices 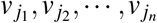, and there exists a unique path 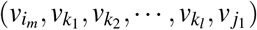 from 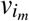 to 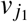 in *G*, and that *m* + *n* + *l* ≥ 3, we then add a phasing path 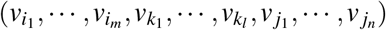 to *H*. In the following, we shall equivalently represent each phasing path with *k* vertices as a consecutive list of (*k* − 1) edges. Different reads or paired-end reads might produce the same phasing path. For each phasing path *h* ∈ *H*, we use *g*(*h*) to record the number of such reads or paired-end reads that produce *h*.

Based on *G, w* and *H*, we compute a set *P* of *s*-*t* paths of *G* and associate a real-value *f*(*p*) for every path *p* ∈ *P*. Each path *p* ∈ *P* implies an expressed transcript, and *f*(*p*) estimates the expression abundance of the corresponding transcript. We now design three objectives to guide reconstructing *P* and *f*. First, since each phasing path is constructed from a single read or paired-end reads, which must be sampled from a single transcript, we expect that each phasing path appears as a whole in some reconstructed transcript. Formally, we say a phasing path *h* ∈ *H* is covered by *P*, if there exists an *s*-*t* path *p* ∈ *P* such that *h* is a consecutive subset of edges of *p*. We do not enforce that all phasing paths in *H* must be covered by *P*. This is because there exist false positive edges in the splice graph due to alignment errors or sequencing errors. Our algorithm will try to identify and remove these false positive edges. Except these phasing paths with false positive edges, we do require that all other phasing paths in *H* are covered by *P*. Second, for each edge *e* ∈ *E* we expect that the superposition of the abundances of the inferred *s*-*t* paths passing through *e*, i.e., 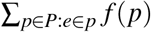, is as close to its observed read coverage *w*(*e*) as possible. Therefore, the second objective is to minimize the deviation between these two quantities, defined as

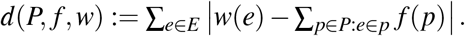

Third, following the principle of parsimony, we expect to use a smaller set of *s*-*t* paths to explain *G, w* and *H*. That is, the third objective is to minimize |*P*|.

Combining all the three objectives, we informally describe the task of transcript assembly as follows.

#### Problem 1 (Transcript Assembly)

*Given G, w and H, compute a set of s-t paths P of G and abundance f(p) for each p* ∈ *P, such that P covers all phasing paths in H (except those with false positive edges), and that both d*(*P, f, w*) *and* |*P*| *are as small as possible*.

### Algorithm

Our algorithm employs an iterative strategy to gradually decompose the splice graph into *s*-*t* paths while achieving the three objectives above. Specifically, we divide all vertices into three types based on the influence of the phasing paths on each vertex, and design different subroutines to decompose each type of vertices. In each iteration, our algorithm decomposes a single vertex so as to either locally minimize the deviation *d*(*P, f, w*), or minimize the number of reconstructed paths |*P*|, while preserving all phasing paths in *H*. Our algorithm can guarantee that, except the phasing paths containing false positive edges, all other phasing paths can be covered by the final set of *s*-*t* paths. This property is achieved by enforcing all three subroutines to keep the invariant that after each iteration every phasing path can be covered by some *s*-*t* in the current splice graph.

We say a vertex *v* ∈ *V* \ {*s*,*t*} is trivial, if its in-degree is 1, or its out-degree is 1; otherwise we say *v* is nontrivial. Intuitively, there is a unique way to decompose a trivial vertex, while there might be multiple ways to decompose a nontrivial vertex. For those nontrivial vertices, we introduce a data structure to further classify them into two types based on the influence of phasing paths on them. For any nontrivial vertex *v*, we build a bipartite graph *G_v_* = (*S_v_* ⋃ *T_v_, E_v_*), in which its vertices (*S_v_* ∪ *T_v_*) correspond to edges in *G*, while its edges (*E_v_*) describe whether the corresponding two edges in *G* are connected by some phasing path in *H*. Formally, let *S_v_* be the set of edges that point to *v*, and let *T_v_* be the set of edges that leave *v*, i.e., *S_v_* = {*e* ∈ *E* | *e* = (*u, v*)} and *T_v_* = {*e* ∈ *E* | *e* = (*v, w*)}. For each pair of edges *e* ∈ *S_v_* and e′ ∈ *T_v_*, we add an edge (*e, e*′) to *E_v_* if there exists a phasing path *h* ∈ *H* such that (*e, e*′) is a consecutive pair in *h*. We say a nontrivial vertex *v* is unsplittable if all elements of *S_v_*, or all elements of *T_v_*, are in the same connected component of *G_v_* (Figure 3(a,b)); otherwise we say *v* is splittable (Figure 5(a,b)). In the following, we design different subroutines to decompose unsplittable vertices, splittable vertices, and trivial vertices.

**Figure 3:**
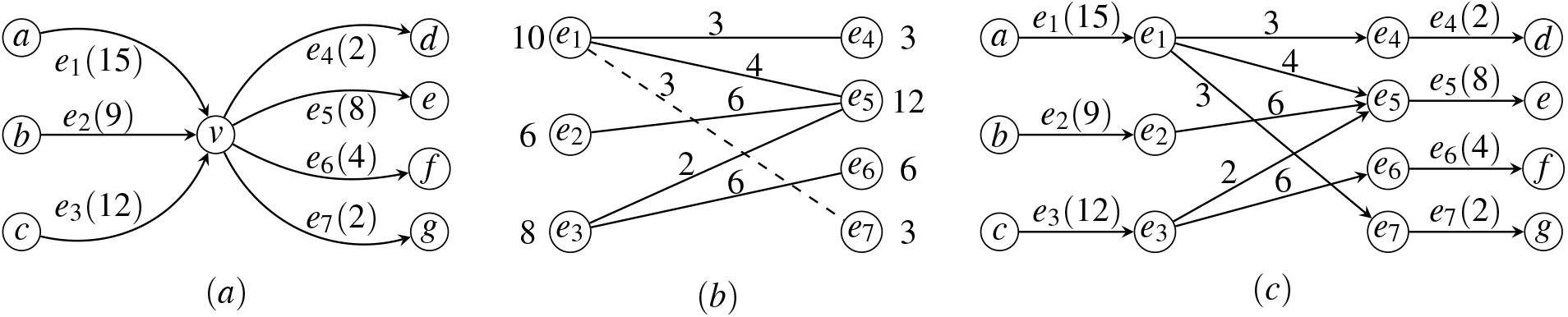
Example of decomposing an unsplittable vertex. (**a**) Subgraph associated with vertex v. The weight of each edge is shown in the parenthesis. Assume that phasing paths contain (*e*_1_, *e*_4_), (*e*_1_, *e*_5_), (*e*_2_, *e*_5_), (*e*_3_, e_5_) and (*e*_3_, *e*_6_). (**b**) Bipartite graph *G_v_* (without the dashed edge), and the extended bipartite graph *G_v_* (with the dashed edge). The balanced weights are next to the vertices. The weights given by the optimal solution of the linear programming are next to edges. (**c**) Updated subgraph after decomposing *v*.

#### Decomposing Unsplittable Vertices

We now describe the subroutine to decompose an unsplittable vertex *v* (Figure 3). The aim of this subroutine is to replace *v* as a set of trivial vertices so as to locally minimize *d*(*P, f, w*) and also preserve all phasing paths.

The first step is to balance *v* by computing new weights 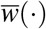 for adjacent edges of *v* (i.e., *S_v_* ∪ *T_v_*). Specifically, for any subset *E*_1_⊂ *E* we define 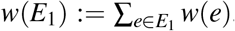. Let 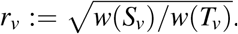. Then we set 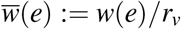 for any edge *e* ∈ *S_v_*, and set 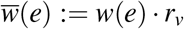 for any edge *e* ∈ *T_v_*. Similarly, we define 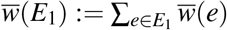 for any subset *E*_1_ ⊂ *E*. Clearly, we have that 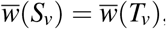, i.e., after balancing the sum of weights of all in-edges of *v* equals that of all out-edges of *v*.

The second step of the subroutine is to build the extended bipartite graph 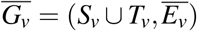.The goal of this extension is to connect edges with no phasing paths to the most likely preceding or succeeding edge. Specifically, let 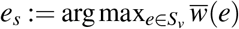 and 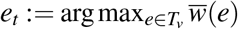 be the edges that have the largest balanced weights in *S_v_* and *T_v_*, respectively. Let 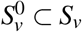 and 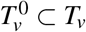 be the set of edges that have total degree of 0 in *G_v_*. We then set 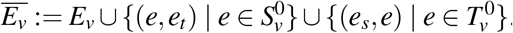. See Figure 3(b) for an example.

The third step of the subroutine is to assign weights for all edges 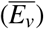 in the extended bipartite graph 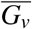 so as to locally minimize the deviation *w.r.t*. 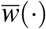, i.e., *d*(*P, f, w*). We formulate it as a linear programming problem. For each edge 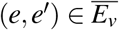 (recall that each edge in 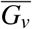 corresponds to a pair of edges in the splice graph *G*), we have a variable *x_e,e′_* to indicate the desired weight of edge (*e, e′*). For each vertex *e* ∈ *S_v_* ∪ *T_v_* (recall that each vertex in 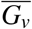 corresponds to an edge in *G*) we add a variable *y_e_* to indicate the deviation between its balanced weight 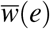 and the sum of the weights of all edges that are adjacent to vertex *e* in 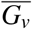. Formally, we have the following constraints:

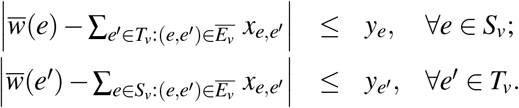

The objective function of the linear programming instance is taken to be:

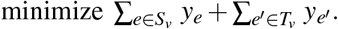

In most cases, when 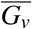 is not a tree, i.e., it contains a cycle (see Figure 4), the above linear programming has multiple optimal solutions. We use the abundance information of the phasing paths stored in *g*(·) to reassign weights while keeping the optimal deviation *w.r.t*. 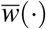. For each edge 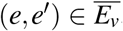, we denote by *g*(*e, e*′) the number of reads or paired-end reads that continuously go through *e* and *e*′, which can be computed as 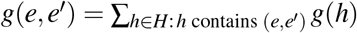. Our goal is then to reassign the weights for edges in 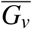 so as to keep the above minimal deviation *w.r.t*. 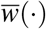 but to minimize the deviation *w.r.t. g*(·, ·).

**Figure 4:**
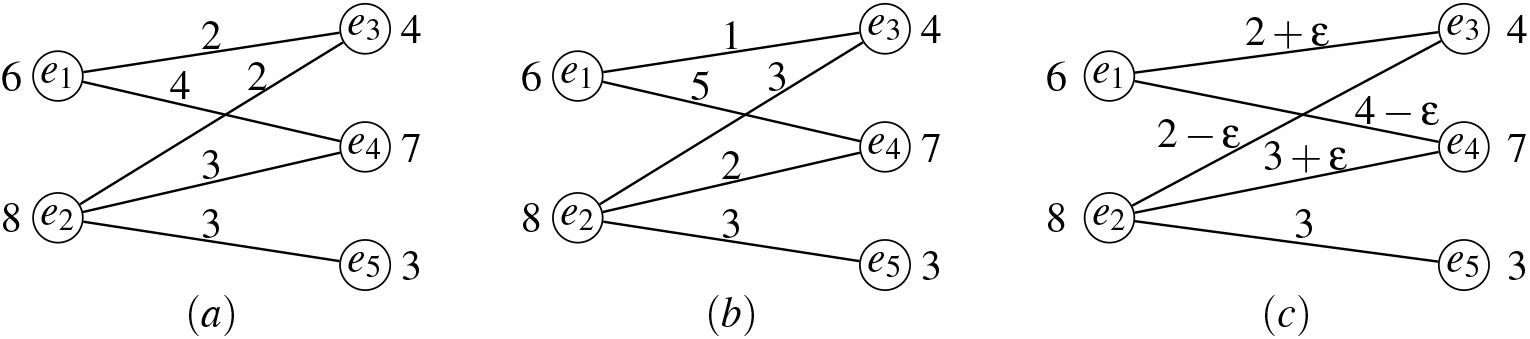
Example with multiple optimal solutions when the extended bipartite graph contains cycles. The balanced weights, i.e., 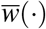, are next to the vertices. (**a, b**) Two optimal solutions with deviation of 0 *w.r.t*. 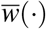. (**c**) When −2 < ε < 2, the solution is always optimal.

We formulate this problem as another linear programming instance. Specifically, let 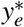 and 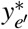, *e* ∈ *S_v_* and *e*′ ∈ *T_v_*, be the optimal solution of first linear programming instance (thus 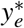 and 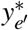 are constants rather than variables in the second linear programming problem). Similar to the first linear programming problem, we use variables *x_e,e_*_′_ to indicate the weight of edge (*e, e*′) in 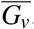. We use the following constraints to guarantee that the optimal weights have the same deviation *w.r.t*. 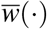 as the first linear programming solution:

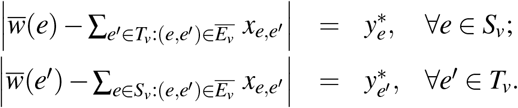

The objective function of this linear programming instance is then taken to minimize the sum of the deviation of weights *w.r.t. g*(·, ·):

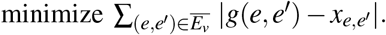

We assign the weights for edges in 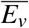 to be the optimal value of *x_e,e_*_′_ of the second linear program. Note that if the first linear program has the unique optimal solution, then both linear programs will have the same optimal solution.

Finally, we update splice graph *G* by replacing *v* with 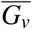 (see Figure 3(**c**)); we denote by *G*′ the updated splice graph. For the edges in 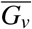 that added to *G*′, we maintain the information that they are artificially added edges and thus do not correspond to any edge in *G*. This information will be used after decomposing all vertices to backtrace the paths with respect to the original splice graph (see line 6 of Algorithm 1). For example, if 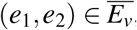, then the path (…, *e*_1_, (*e*_1_, *e*_2_), *e*_2_, …) in *G*′ corresponds to the path (…, *e*_1_, *e*_2_, …) in *G*. We then update *H*; we denote by *H*′ the updated set of phasing paths. For any phasing path *h* ∈ *H*, if *h* contains a pair of continuous edges *e* and *e*′ in *G* such that 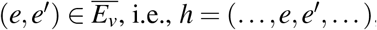, then *h* will become *h*′ = (…, *e*, (*e, e*′), *e*′; …) ∈ *H*′.

This subroutine preserves all phasing paths in *H*, i.e., every phasing path *h* ∈ *H* is still covered by some *s*-*t* path *p*′ of *G*′ (i.e, if we transform *p*′ of *G*′ into the corresponding path *p* of *G* through removing the artificially added edges in *p*′ (if any), then *p* covers *h*). This is true because according to our construction of *G*′ and *H*′ there is a one-to-one correspondence between *H* and *H*′ and every phasing path in *H*′ is covered by some path in *G*′.

To choose which unsplittable vertex *v* to apply the above transformation to, we define

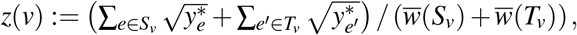

and select a vertex *v* that minimizes *z*(*v*) to decompose (see line 3 of Algorithm 1).

### Decomposing Splittable Vertices

We now describe the subroutine to decompose a splittable vertex *v* (Figure 5). The aim of this subroutine is to reduce |*P*| while preserving all phasing paths. Since *P* is not explicitly available until we have finished decomposing all vertices, we use *U* := |*E*| − |*V*| + 2 to approximate |*P*|. It has been proved that ∪ is an upper bound of |*P*| in the flow decomposition scenario: for a given flow at most *U* paths are required to decompose this flow [32, 33]. Following this approximation, in order to reduce |*P*|, our subroutine to decompose a splittable *v* will increase the number of vertices or decrease the number of edges, while at the same time preserving all phasing paths. Splittable vertices will typically be replaced by two new vertices.

**Figure 5:**
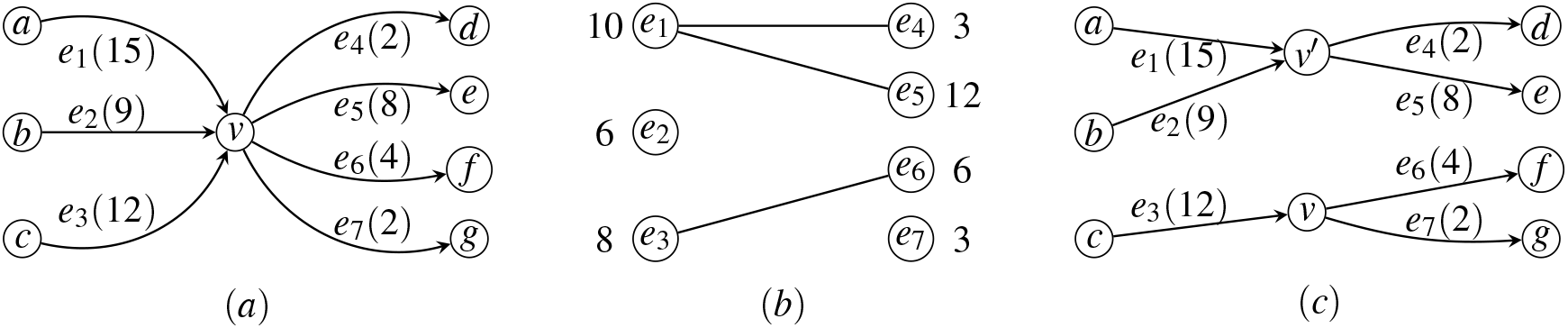
Example of decomposing a splittable vertex. (**a**) Subgraph associated with *v*. Assume that phasing paths contain (*e*_1_, *e*_4_), (*e*_1_, *e*_5_) and (*e*_3_, *e*_6_). (**b**) Bipartite graph *G_v_* with balanced weights next to vertices. Optimal decomposition gives 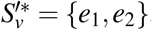, 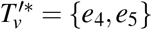. (**c**) Updated subgraph after decomposing *v*.

The first step of this subroutine is also to balance *v* by computing 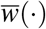 for edges in *S_v_* ∪ *T_v_* following the same procedure as in decomposing unsplittable vertices. The second step is to split *v* into two vertices so as to keep all phasing paths and to minimize the balanced weight discrepancy (i.e., to make each of the two new vertices as balanced as possible). Formally, we seek *S_v_*′ ⊂ *S_v_* and *T_v_*′ ⊂ *T_v_, S_v_*′ ∪ *T_v_*′ ≠ Ø and *S_v_*′ ∪ *T_v_*′ ≠ *S_v_* ∪ *T_v_*, such that for each (*e, e*′) ∈ *E_v_*, either *e* ∈ *S_v_*′ and *e*′ ∈ *T_v_*′, or 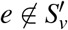 and 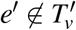, and that 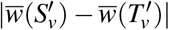 is minimized. Intuitively, this formulation forces that two edges in some phasing path must be adjacent after splitting, and thus all phasing paths can be preserved.

The above problem can be equivalently transformed into the subset-sum problem. Let 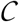 be the set of all connected components of *Gv*. We define 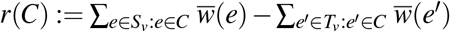, for any 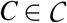. Then the above problem is equivalent to computing a nonempty and strict subset of 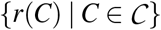 such that the sum of all elements of this subset is closest to 0. In our implementation, we use the existing pseudo-polynomial time dynamic programming algorithm to solve it.

Let 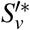 and 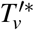 be the optimal subsets returned by the above algorithm. We then update splice graph *G* through performing the following procedure to decompose *v*. We denote the updated splice graph as *G*′ (see Figure 5). Vertex *v* will be split into two vertices by adding another vertex *v*′ to *G*′. Edges in 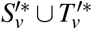 will be detached from *v* and attached to *v*′. Notice that the weights for edges in *S_v_* ∪ *T_v_* are kept unchanged as *w*(·), i.e., the balanced weight 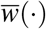 is only used to compute 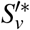 and 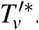.

It could be the case that 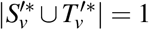 (or symmetrically, 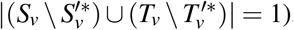, i.e., the new vertex *v*′ will have either in-degree of 0 and out-degree or 1, or out-degree of 0 and in-degree of 1. In this case, the above procedure of decomposing *v* will degenerate into removing this edge from *G*, instead of splitting *v* into two vertices. If this is the case, it indicates that this particular edge is more likely to be a false positive edge. In other words, this procedure can be used to naturally remove false positive edges in the splice graph. For this case, we remove the appearance of this false positive edge for all phasing paths in *H*.

Notice that in either the general case of splitting *v* into two vertices, or the degenerate case of removing one edge from *G*, after decomposing splittable vertex *v*, we have that *U* will be reduced by 1.

For the degenerated case of removing one edge from *G*, these spanning paths that contain this edge shall be not covered by *G*′. For the usual case of splitting vertex this subroutine keeps all phasing paths *H* unchanged. Finally, we define

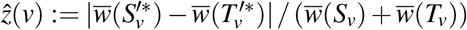

as the measurement to decide which splittable vertex to decompose (see line 4 of Algorithm 1).

#### Decomposing Trivial Vertices

There is a unique way to decompose a trivial vertex. Let *v* ∈ *V* be a trivial vertex. Again, let *S_v_* be the set of edges that point to *v*, and let *T_v_* be the set of edges that leave *v*. Without loss of generality, we assume that the in-degree of *v* is 1; let *e* = (*u, v*) be the only in-edge of *v*, i.e., *S_v_* = {*e* = (*u, v*)}. We denote by *G*′ the updated splice graph after decomposing *v*. The construction of *G*′ from *G* is to remove edge *e* from *G*, and merge *u* and *v* as a single vertex *v* (Figure 6). For each edge in *e*′ ∈ *T_v_*, we maintain the information that *e*′ is preceded by an extra edge *e*, i.e., for *e*′ in *G* we label it as *ee*′ in *G*′. When we retrieve the paths *w.r.t*. the original splice graph (line 6 of Algorithm 1), *ee*′ in *G*′ will be expanded as a pair of edges (*e, e*′) in *G*. We then update the phasing paths *H*; we denote by *H*′ the updated set of phasing paths. Consider two cases of a phasing path *h* ∈ *H* that contains *e*. If *e* is the last edge of *h*, i.e., *h* = (…, *e*_1_, *e*), then we simply remove *e* from *h*, i.e., it becomes *h*′ = (…, *e*_1_) ∈ *H*′. Otherwise, if *e* is not the last element of *h*, i.e., *h* = (…, *e, e*_1_,…), we replace *e* and *e*_1_ as the edge *ee*_1_ in *G*′, i.e., it becomes *h*′ = (…, *ee*_1_,…) ∈ *H*′.

**Figure 6:**
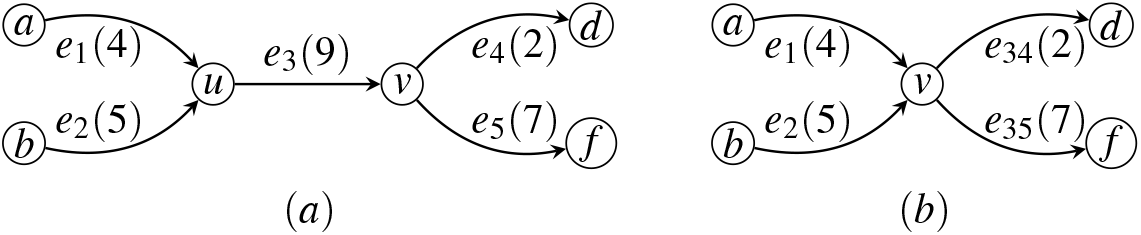
Illustration of decomposing trivial vertices *v*. (**a**) Subgraph before decomposing *v*. (**b**) Subgraph after decomposing *v*. Notice that we maintain the information that *e*_4_ and *e*_5_ are preceded by *e*_3_ by labeling them as *e*_34_ and *e*_35_.

In the complete algorithm (Algorithm 1), we first decompose all nontrivial vertices before decomposing any trivial vertex. In other words, when we use the above subroutine to decompose a trivial vertex, all vertices in the current splice graph are trivial vertices. We now prove that, when the splice graph contains only trivial vertices, our subroutine to decompose trivial vertex also preserves all phasing paths. Again, consider the two cases of a phasing path *h* ∈ *H* that contains *e*. If *h* = (…, *e, e*_1_,…) ∈ *H*, then we have *h*′ = (…, *ee*_1_, …) ∈ *H*′, and *h*′ is covered by *G*′. Since *ee*_1_ is essentially the concatenation of *e* and *e*_1_, we have that *G*′ covers *h*. For the other case that *h* = (…, *e*_1_, *e*) ∈ *H*, we have that *h*′ = (…, *e*_1_) ∈ *H*′. (Although *G*′ covers *h*′, but this alone does not necessarily imply that *G*′ covers *h* any more.) Let *e*_1_ = (*w, u*) and *e* = (*u, v*) in *G*. Since we assume that all vertices in *G* are trivial vertices, in particular *u* is a trivial vertex, we have that in *G*′ all the succeeding edges of *e*_1_ are in the type of *ee*′, where *e*′ ∈ *T_v_* (see Figure 6). In other words, for any path in *G*′ that contains *h*′, the next edge of *e*_1_ in this path must be an concatenated edge with preceding edge of *e*. Hence, we have that *G*′ covers *h*.

We emphasize that if the splice graph contains nontrivial vertex, then decomposing a trivial vertex might not preserve all phasing paths. Figure 7 gives such an example. Thus, it is essential to decompose all nontrivial vertices before decomposing any trivial vertex.

**Figure 7:**
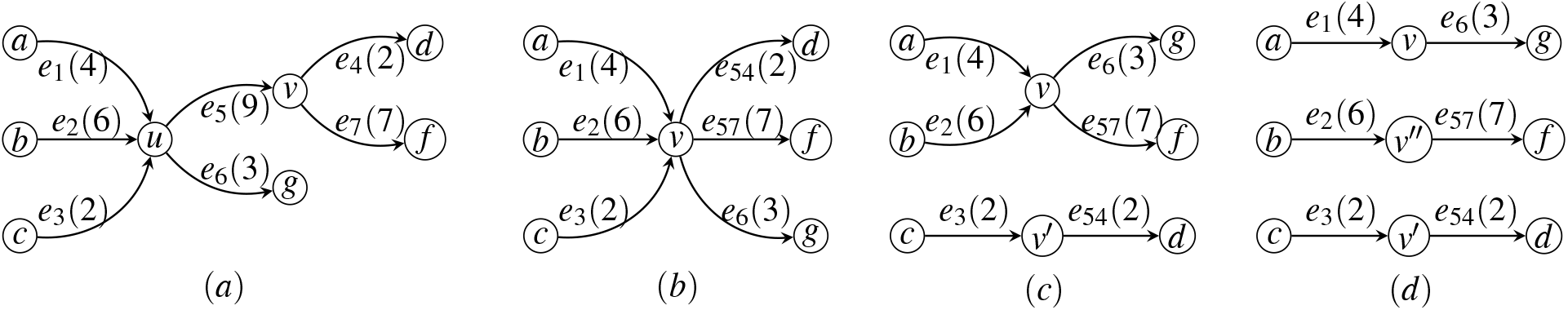
Decomposing a trivial vertex may not preserve all phasing paths if the splice graph contain non-trivial vertices. (**a**) Splice graph *G* with trivial vertex *v* and nontrivial vertex *u*. Assume that we have a single phasing path of *H* = {(*e*_1_, *e*_5_)}. (**b**) Updated splice graph *G*′ after decomposing trivial vertex *v*. Notice that now we have *H*′ = {(*e*_1_)}. Since a phasing path with a single edge is not informative, we actually have that *H*′ = Ø. (**c,d**) The following decomposition of *G*′ by applying the subroutine for splittable vertices. Notice that in the final three *s*-*t* paths, none of them covers (*e*_1_, *e*_5_).

#### Complete Algorithm

Our complete algorithm for Problem 1 is to iteratively decompose vertices by applying the above three subroutines, until finally the splice graph becomes a set of *s*-*t* paths. The complete algorithm is in Algorithm 1. Notice that when the Algorithm 1 reaches line 5, all vertices must be trivial vertices. Among nontrivial vertices, we further give priority to unsplittable ones, since their decomposition is fully determined by phasing paths.

##### Algorithm 1: Heuristic for Problem 1

**Input:** *G, w, H* and *g*

**Output:** *P* and *f*

1. Let *V*_1_ ⊂ *V* \ {*s*,*t*} be the set of unsplittable vertices.

2. Let *V*_2_ ⊂ V\ {*s*,*t*} be the set of splittable vertices.

3. **If** *V*_1_ ≠ Ø, compute 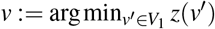, decompose *v* by updating *G, w* and *H*, and **goto** step 1.

4. **If** *V*_2_ ≠ Ø, compute 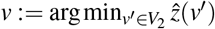, decompose *v* by updating *G*, and **goto** step 1.

5. Arbitrarily choose a (trivial) vertex *v* ∈ *V*\ {*s*,*t*}, decompose *v* by updating *G* and *H*, and **goto** step 1.

6. For all the *s*-*t* edges of *G*, recover the original *s*-*t* paths as *P*; set *f* as the corresponding weights of the edges.

---

**Supplementary Figure 1:**
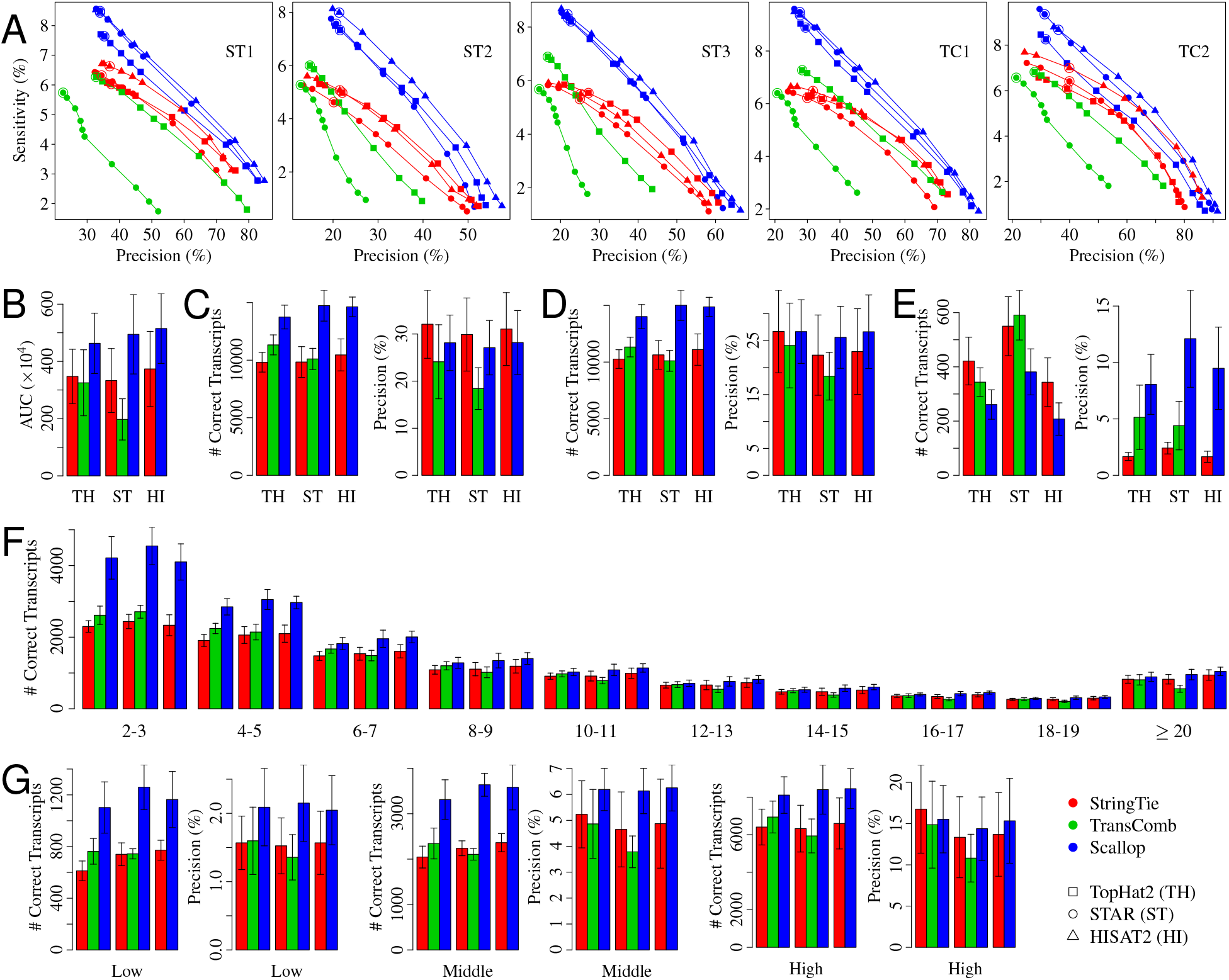
Comparison of the three methods (StringTie, TransComb, and Scallop) over the 5 training samples. (A) The precision-sensitivity curves for multi-exon transcripts. Each curve connects 10 points, corresponding to the 10 different minimum coverage thresholds {0, 1, 2.5, 5, 7.5, 10, 25, 50, 75, 100}; the default value of this parameter is circled. (B) The average AUC (area under the precision-sensitivity curve). The three groups of bars correspond to TopHat2, STAR, and HISAT2 alignments, respectively (the same for other panels). The error bars show the standard deviation over the 5 samples (the same for other panels). (C) The average sensitivity and precision of multi-exon transcripts for methods running with default parameters. (D) The average sensitivity and precision of multi-exon transcripts for methods running with minimum coverage set to 0. (E) The average sensitivity and precision of single-exon transcripts for methods running with default parameters. (F) The average number of correct transcripts with different number of exons for methods running with minimum coverage set to 0. (G) The average sensitivity and precision of multi-exon transcripts with each subset of transcripts (corresponding to low, middle, and high expression level) as ground truth for methods running with minimum coverage set to 0.

**Supplementary Figure 2:**
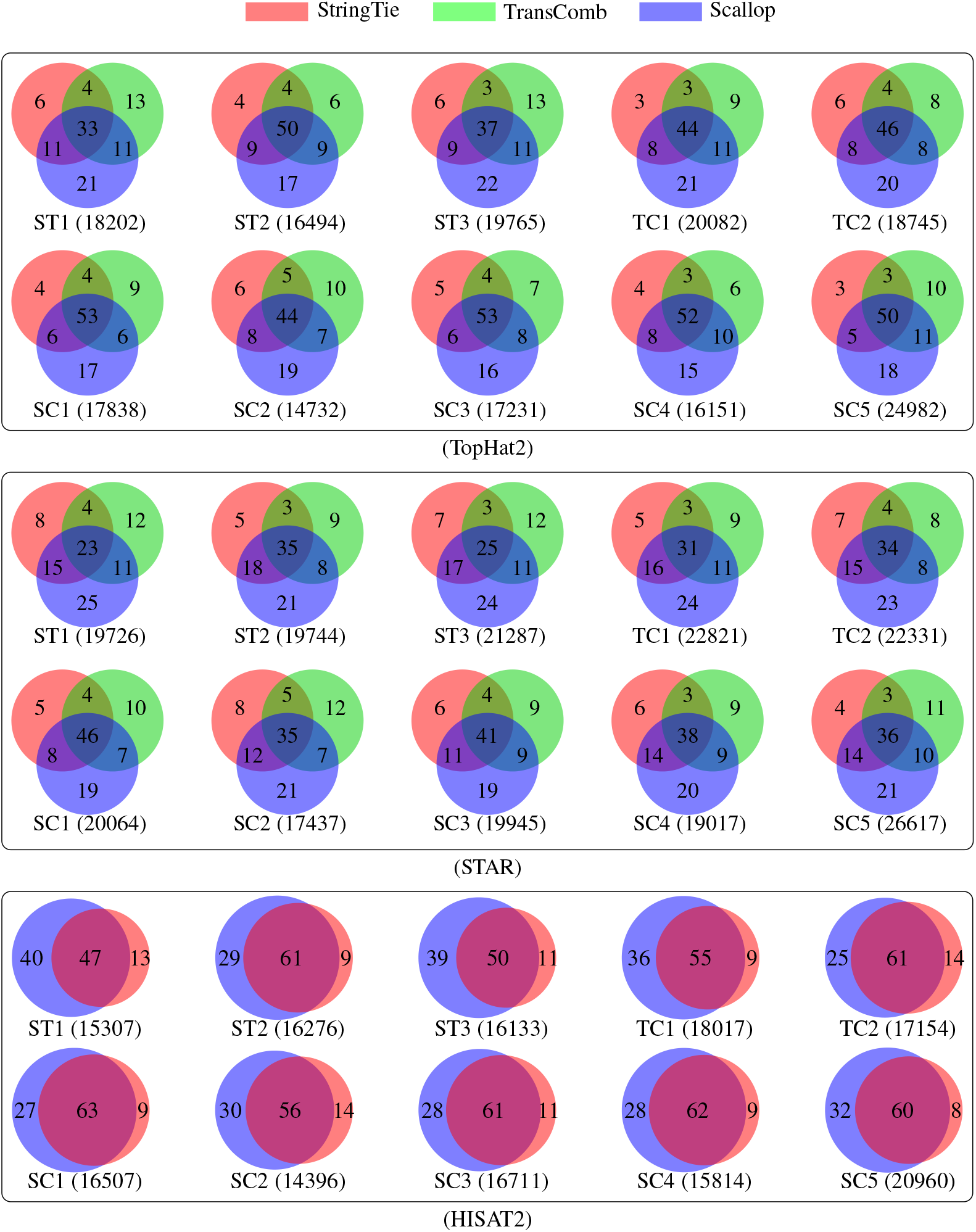
Correlation among different assemblers. For each Venn diagram, the number in the parenthesis below gives the number of correct transcripts in the union of all three assemblers. The numbers inside the Venn diagram gives the percentage of the correct transcripts in the corresponding subset with respect to the union. The three assemblers are very diverse from each other. Specifically, the ratio between the number of correct transcripts shared by all the three assemblers and that in the union of them is 43.1% and 31.0% for TopHat2 and STAR alignments, respectively. With HISAT2 alignments, the ratio between the number of correct transcripts shared by StringTie and Scallop and that in the union of them is 57.5%.

**Supplementary Figure 3:**
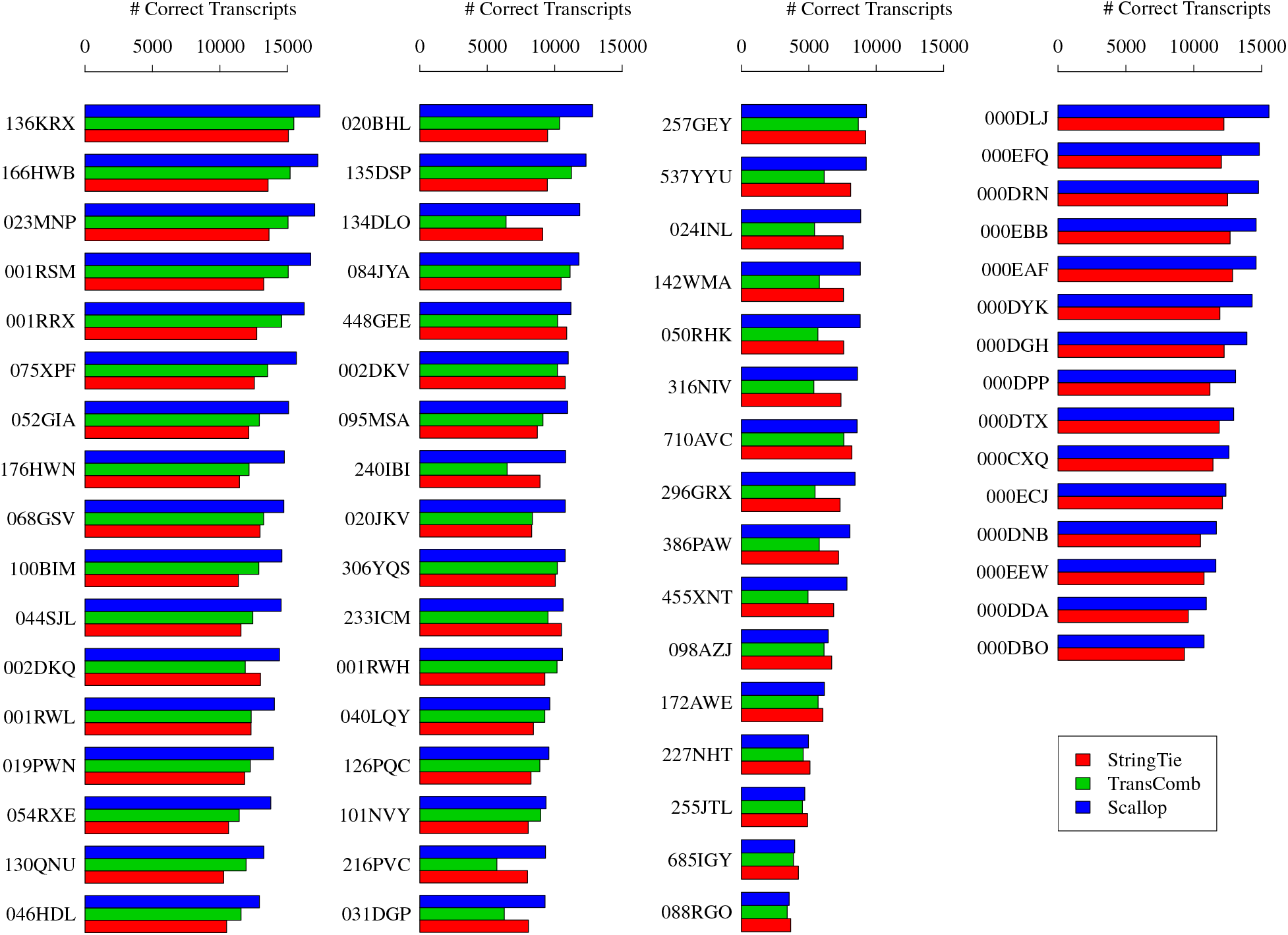
Comparison of the adjusted sensitivity (shown as the number of correct transcripts) of multi-exon transcripts for methods (StringTie, TransComb, and Scallop) running with their default parameter settings. The experiment uses 50 strand-specific samples (leftmost three columns of this figure) and 15 non-strand-specific samples (the rightmost column of this figure). Read alignments for these samples were downloaded from ENCODE project (2013–present). For each sample, we mark its (partial) ID in ENCODE on the left side. The complete ID adds the prefix “ENCFF”. TransComb fails on the 15 non-strand-specific samples so for them we only compare the results given by Scallop and StringTie.

**Supplementary Figure 4:**
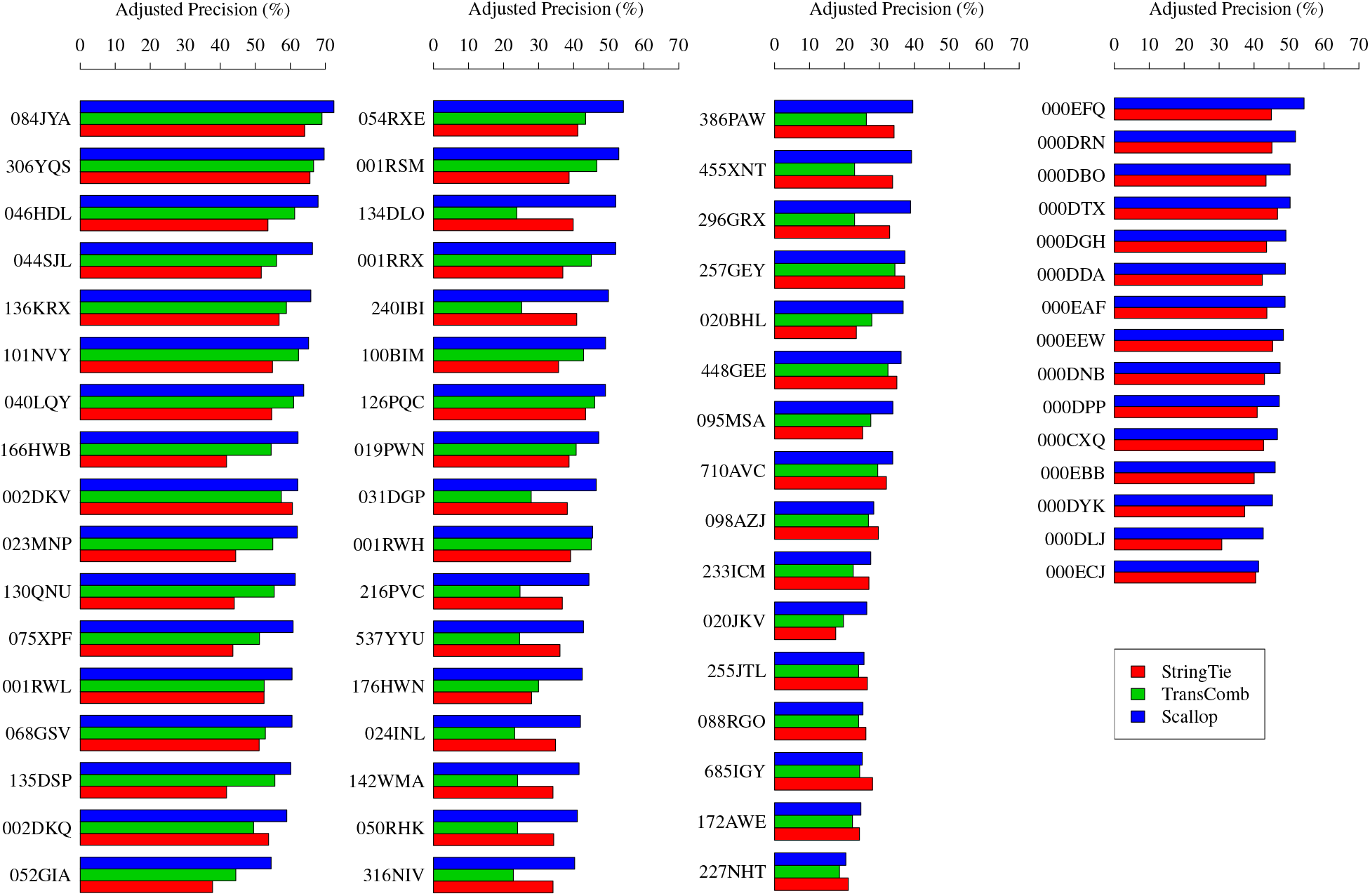
Comparison of the adjusted precision of multi-exon transcripts for methods (StringTie, TransComb, and Scallop) running with their default parameter settings. The samples are identical to those in Supplementary Figure 3.

**Supplementary Figure 5:**
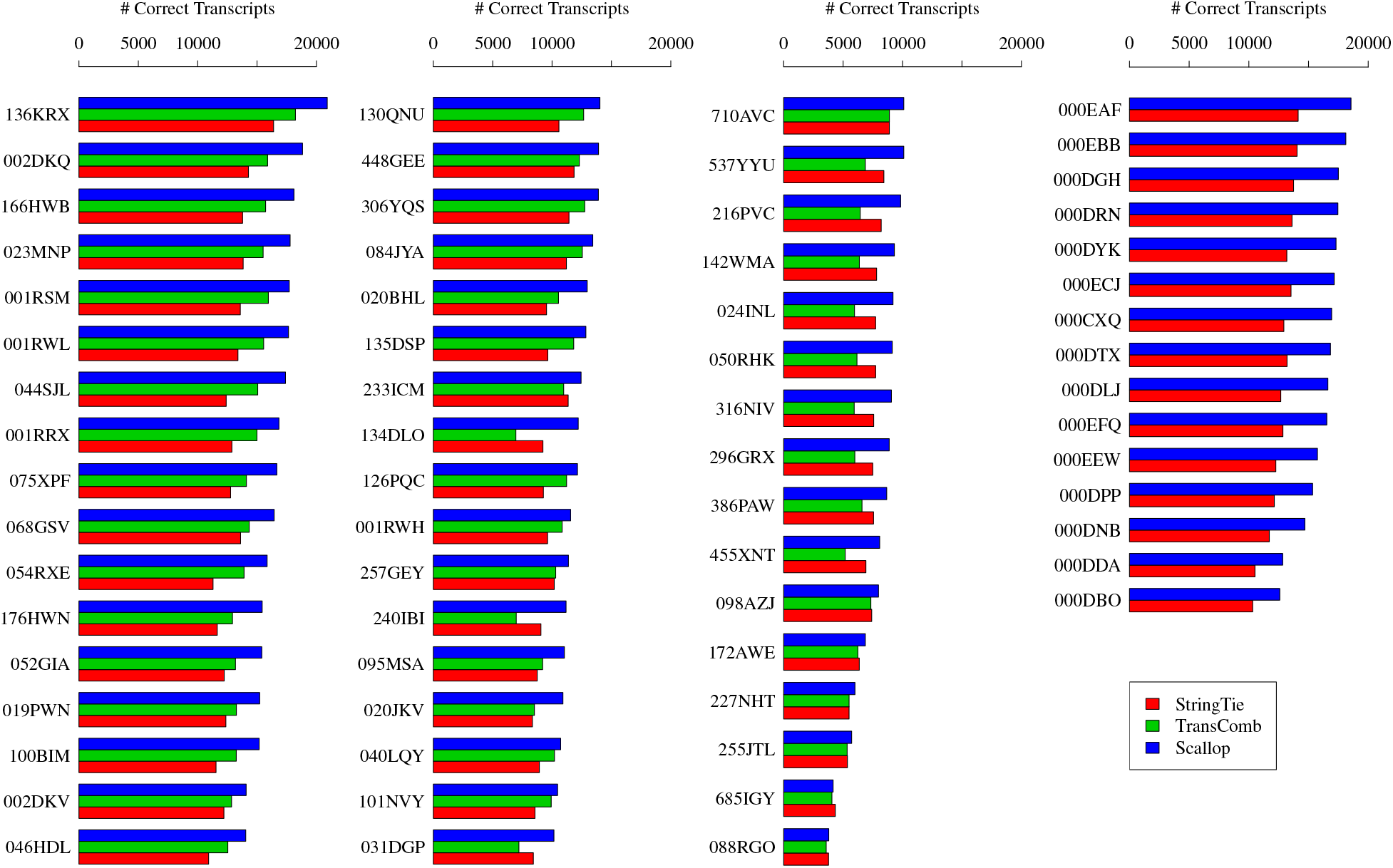
Comparison of the adjusted sensitivity (shown as the number of correct transcripts) of multi-exon transcripts for methods (StringTie, TransComb, and Scallop) running with the minimum coverage threshold set to 0. The samples are identical to those in Supplementary Figure 3. Scallop produces higher adjusted sensitivity than StringTie on 64 out of the 65 samples, and than TransComb on all the 50 strand-specific samples. (TransComb fails on all non-strand-specific samples.) On average over the 50 strand-specific samples, Scallop obtains 24.1% and 18.7% more correct multi-exon transcripts (after adjustment) than StringTie and TransComb, respectively. Averaged over all the 15 non-strand-specific samples, Scallop finds 28.0% more correct multi-exon transcripts (after adjustment) than StringTie.

**Supplementary Figure 6:**
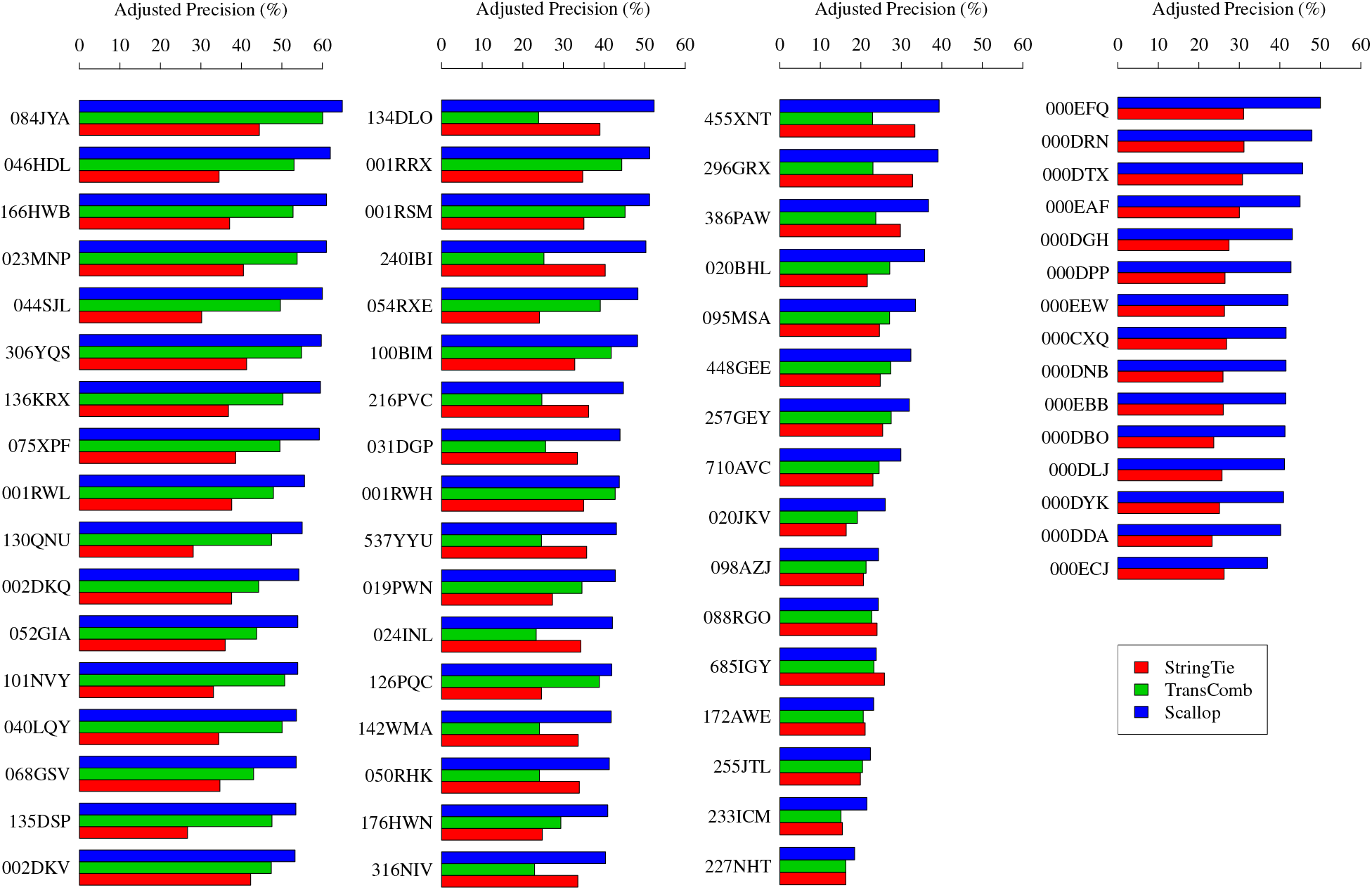
Comparison of the adjusted precision of multi-exon transcripts for the three methods (StringTie, TransComb, and Scallop) running with the minimum coverage threshold set to 0. The samples are identical to those in Supplementary Figure 3. Scallop produces higher adjusted precision than StringTie on 64 out of the 65 samples, and than TransComb on all the 50 strand-specific samples. (TransComb fails on all non-strand-specific samples.) On the 50 strand-specific samples, the average adjusted precision is 30.9%, 34.8%, and 44.1% for StringTie, TransComb, and Scallop, respectively. On the 15 non-strand-specific samples, the average adjusted precision for Scallop is 42.8%, significantly outperforming StringTie at 27.1%.

**Supplementary Figure 7:**
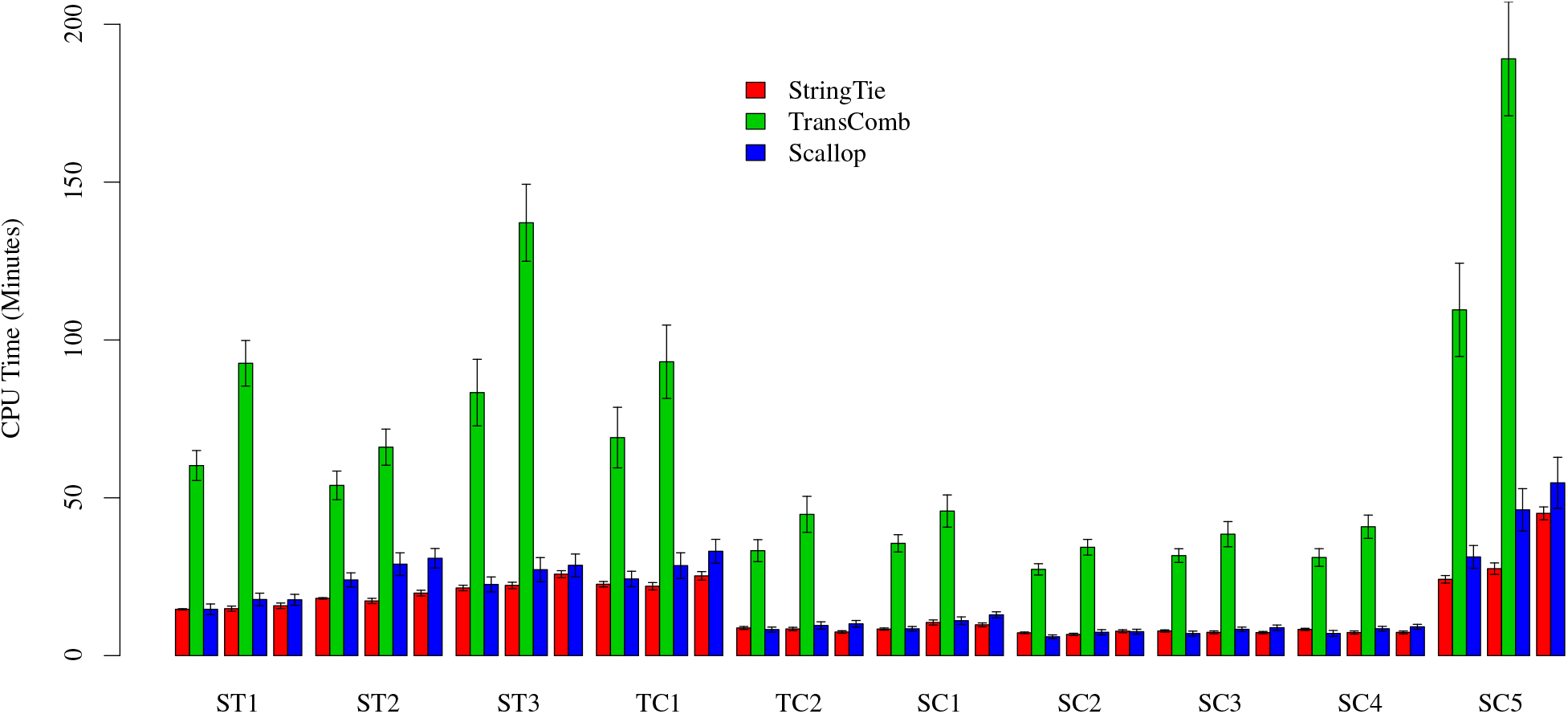
Comparison of the running time (measured as CPU time) of the three assemblers (StringTie, TransComb, and Scallop) on 10 RNA-seq samples. All programs are run with their single-thread mode on a machine with 48 cores and 40GB RAM. The error bars show the standard deviation over the 10 runs with different minimum coverage parameters {0, 1, 2.5, 5, 7.5, 10, 25, 50, 75, 100}.

**Supplementary Table 1:**
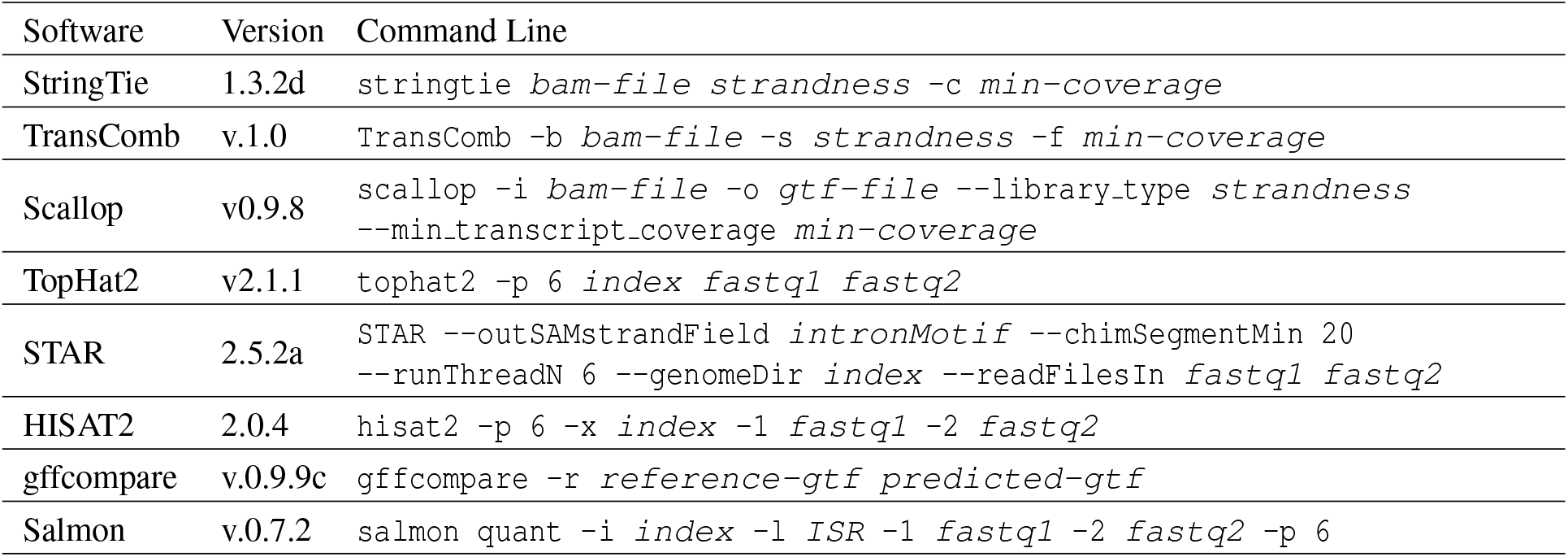
Programs and their versions and arguments used in this paper.

**Supplementary Table 2:** Summary of the 10 RNA-seq samples used in this paper (except samples in Supplementary Figures 3). All these 10 samples are from human, and the sequencing employs paired-end and strand-specific protocols. All datasets are downloaded from ENCODE project (2003–2012).

| ID | SRA Accession | GEO Accession | Chosen By | #Spots | Cell Line | Localization | Length |
| --- | --- | --- | --- | --- | --- | --- | --- |
| ST1 | SRR534319 | GSM981256 | StringTie | 25M | CD20+ | cell | 76 |
| ST2 | SRR534291 | GSM981244 | StringTie | 114M | IMR90 | cytosol | 101 |
| ST3 | SRR545695 | GSM984609 | StringTie | 40M | CD14+ | cell | 76 |
| TC1 | SRR387661 | GSM840137 | TransComb | 125M | K562 | cytosol | 76 |
| TC2 | SRR307911 | GSM758566 | TransComb | 41M | H1-hESC | cell | 76 |
| SC1 | SRR545723 | GSM984621 | Scallop | 147M | HMEpC | cell | 101 |
| SC2 | SRR315323 | GSM765399 | Scallop | 30M | NHEK | nucleus | 76 |
| SC3 | SRR307903 | GSM758562 | Scallop | 36M | BJ | cell | 76 |
| SC4 | SRR315334 | GSM765404 | Scallop | 39M | HeLa-S3 | cytosol | 76 |
| SC5 | SRR534307 | GSM981252 | Scallop | 167M | MCF-7 | cytosol | 101 |

## Reference

[1] M. Pertea, G.M. Pertea, C.M. Antonescu, T.-C. Chang, J.T. Mendell, and S.L. Salzberg. StringTie enables improved reconstruction of a transcriptome from RNA-seq reads. Nat. Biotechnol., 33(3): 290–295, 2015.

[2] J. Liu, T. Yu, T. Jiang, and G. Li. TransComb: genome-guided transcriptome assembly via combing junctions in splicing graphs. Genome Biol., 17(1):213, 2016.

[3] Z. Wang, M. Gerstein, and M. Snyder. RNA-Seq: a revolutionary tool for transcriptomics. Nat. Rev. Genet., 10(1):57–63, 2009.

[4] N.L. Barbosa-Morais, M. Irimia, Q. Pan, H.Y. Xiong, S. Gueroussov, L.J. Lee, et al. The evolutionary landscape of alternative splicing in vertebrate species. Science, 338(6114):1587–1593, 2012.

[5] J.K. Pickrell, J.C. Marioni, A.A. Pai, J.F. Degner, B.E. Engelhardt, E. Nkadori, J.-B. Veyrieras, M. Stephens, Y. Gilad, and J.K. Pritchard. Understanding mechanisms underlying human gene expression variation with RNA sequencing. Nature, 464(7289):768–772, 2010.

[6] C. Trapnell, B.A. Williams, G. Pertea, A. Mortazavi, G. Kwan, M.J. Van Baren, S.L. Salzberg, B.J. Wold, and L. Pachter. Transcript assembly and quantification by RNA-Seq reveals unannotated transcripts and isoform switching during cell differentiation. Nat. Biotechnol., 28(5):511–515, 2010.

[7] M. Guttman, M. Garber, J.Z. Levin, J. Donaghey, J. Robinson, X. Adiconis, L. Fan, et al. Ab initio reconstruction of cell type-specific transcriptomes in mouse reveals the conserved multi-exonic structure of lincrnas. Nat. Biotechnol., 28(5):503–510, 2010.

[8] W. Li, J. Feng, and T. Jiang. IsoLasso: a LASSO regression approach to RNA-Seq based transcriptome assembly. J. Comput. Biol., 18(11):1693–1707, 2011.

[9] J.J. Li, C.-R. Jiang, J.B. Brown, H. Huang, and P.J. Bickel. Sparse linear modeling of next-generation mRNA sequencing (RNA-Seq) data for isoform discovery and abundance estimation. Proc. Natl. Acad. Sci. USA, 108(50):19867–19872, 2011.

[10] Y.-Y. Lin, P. Dao, F. Hach, M. Bakhshi, F. Mo, A. Lapuk, C. Collins, and S.C. Sahinalp. CLIIQ: Accurate comparative detection and quantification of expressed isoforms in a population. In Proc. 12th Workshop Algs. in Bioinf. (WABI’12), volume 7534 of Lecture Notes in Comp. Sci., pages 178–189, 2012.

[11] W. Li and T. Jiang. Transcriptome assembly and isoform expression level estimation from biased RNA-Seq reads. Bioinformatics, 28(22):2914–2921, 2012.

[12] J. Behr, A. Kahles, Y. Zhong, V.T. Sreedharan, P. Drewe, and G. Rätsch. MITIE: Simultaneous RNA-Seq-based transcript identification and quantification in multiple samples. Bioinformatics, 29(20): 2529–2538, 2013.

[13] A.M. Mezlini, E.J.M. Smith, M. Fiume, O. Buske, G.L. Savich, et al. iReckon: Simultaneous isoform discovery and abundance estimation from RNA-seq data. Genome Res., 23(3):519–529, 2013.

[14] A.I. Tomescu, A. Kuosmanen, R. Rizzi, and V. Mäkinen. A novel min-cost flow method for estimating transcript expression with RNA-Seq. BMC Bioinformatics, 14(5):1, 2013.

[15] L. Maretty, J.A. Sibbesen, and A. Krogh. Bayesian transcriptome assembly. Genome Biol., 15(10):1, 2014.

[16] S. Canzar, S. Andreotti, D. Weese, K. Reinert, and G.W. Klau. CIDANE: comprehensive isoform discovery and abundance estimation. Genome Biol., 17(1):16, 2016.

[17] D. Kim, G. Pertea, C. Trapnell, H. Pimentel, R. Kelley, S.L. Salzberg, et al. TopHat2: accurate alignment of transcriptomes in the presence of insertions, deletions and gene fusions. Genome Biol., 14(4):R36, 2013.

[18] K.F. Au, H. Jiang, L. Lin, Y. Xing, and W.H. Wong. Detection of splice junctions from paired-end RNA-seq data by SpliceMap. Nucleic Acids Res., 38(14):4570–4578, 2010.

[19] A. Dobin, C.A. Davis, F. Schlesinger, J. Drenkow, C. Zaleski, S. Jha, P. Batut, M. Chaisson, and T.R. Gingeras. STAR: ultrafast universal RNA-seq aligner. Bioinformatics, 29(1):15–21, 2013.

[20] D. Kim, B. Langmead, and S.L. Salzberg. HISAT: a fast spliced aligner with low memory requirements. Nat. Methods, 12(4):357–360, 2015.

[21] J. Liu, G. Li, Z. Chang, T. Yu, B. Liu, R. McMullen, P. Chen, and X. Huang. BinPacker: Packing-based de novo transcriptome assembly from RNA-seq data. PLoS Comput. Biol, 12(2):e1004772, 2016.

[22] G. Robertson, J. Schein, R. Chiu, R. Corbett, M. Field, S.D. Jackman, K. Mungall, S. Lee, H.M. Okada, J.Q. Qian, et al. De novo assembly and analysis of RNA-seq data. Nat. Methods, 7(11):909–912, 2010.

[23] J. Martin, V.M. Bruno, Z. Fang, X. Meng, M. Blow, T. Zhang, G. Sherlock, M. Snyder, and Z. Wang. Rnnotator: an automated de novo transcriptome assembly pipeline from stranded RNA-Seq reads. BMC Genomics, 11(1):1, 2010.

[24] M.G. Grabherr, B.J. Haas, M. Yassour, J.Z. Levin, D.A. Thompson, et al. Trinity: reconstructing a full-length transcriptome without a genome from RNA-Seq data. Nat. Biotechnol., 29(7):644, 2011.

[25] Y. Xie, G. Wu, J. Tang, R. Luo, J. Patterson, S. Liu, et al. SOAPdenovo-Trans: de novo transcriptome assembly with short RNA-Seq reads. Bioinformatics, 30(12):1660–1666, 2014.

[26] D.R. Zerbino and E. Birney. Velvet: algorithms for de novo short read assembly using de Bruijn graphs. Genome Res., 18(5):821–829, 2008.

[27] M.H. Schulz, D.R. Zerbino, M. Vingron, and E. Birney. Oases: robust de novo RNA-seq assembly across the dynamic range of expression levels. Bioinformatics, 28(8):1086–1092, 2012.

[28] Y. Peng, H.C.M. Leung, S.-M. Yiu, M.-J. Lv, X.-G. Zhu, and F.Y.L. Chin. IDBA-tran: a more robust de novo de Bruijn graph assembler for transcriptomes with uneven expression levels. Bioinformatics, 29(13):i326–i334, 2013.

[29] T. Steijger, J.F. Abril, P.G. Engström, F. Kokocinski, T.J. Hubbard, et al. Assessment of transcript reconstruction methods for RNA-seq. Nat. Methods, 10(12):1177–1184, 2013.

[30] K.E. Hayer, A. Pizarro, N.F. Lahens, J.B. Hogenesch, and G.R. Grant. Benchmark analysis of algorithms for determining and quantifying full-length mRNA splice forms from RNA-seq data. Bioinformatics, 31(24):3938–3945, 2015.

[31] R. Patro, G. Duggal, M.I. Love, R.A. Irizarry, and C. Kingsford. Salmon provides fast and bias-aware quantification of transcript expression. Nat. Methods, 2017. doi:10.1038/nmeth.4197.

## Reference

[32] B. Vatinlen, F. Chauvet, P. Chrétienne, and P. Mahey. Simple bounds and greedy algorithms for decomposing a flow into a minimal set of paths. Eur. J. Oper. Res., 185(3):1390–1401, 2008.

[33] M. Shao and C. Kingsford. Efficient heuristic for decomposing a flow with minimum number of paths. bioRxiv, page 087759, 2016. doi:10.1101/087759.

